# netome: a computational framework for metabolite profiling and omics network analysis

**DOI:** 10.1101/443903

**Authors:** Ali Rahnavard, Daniel Hitchcock, Julian Avila Pacheco, Amy Deik, Courtney Dennis, Sarah Jeanfavre, Kerry Pierce, Kevin Bullock, Zach Costliow, Clary B. Clish

## Abstract

**Summary:** Advances in metabolomics technologies have enabled comprehensive analyses of associations between metabolites and human disease and have provided a means to study biochemical pathways and processes in detail using model systems. Liquid chromatography tandem mass spectrometry (LC-MS) is an analytical technique commonly used by metabolomics labs to measure hundreds of metabolites of known identity and thousands of “peaks” from yet to be identified compounds that are tracked by their measured masses and chromatographic retention times. *netome* is a computational framework that provides tools for analyzing processed LC-MS data. In this framework, we develop and provide various computational resources including individual software modules to inspect and adjust trends in raw data, align unknown peaks between separately acquired data sets, and to remove redundancies in nontargeted LC-MS data arising from multiple ionization products of a single metabolite. These tools are deployed through computing resources such as web servers and virtual machines with detailed documentation in order to support researchers.

**Availability and implementation:** *netome* is publicly available with extensive documentation and support via issue tracker at https://broadinstitute.github.io/netome under the MIT license. *netome* includes a set of computational methods that have been designed to execute quality control and post-raw data processing tasks for metabolomics data (e.g. scaling and clustering metabolite abundances), as well as statistical association testing in a network manner (e.g., testing relationship between metabolites and microbes). Each individual tool is available with source code, workshop-oriented documentation which includes instructions for installation and using tools with demonstration examples, and a web server with all services. We also provide a complete image of the *netome* package with all pre-installed dependencies and support for Google Compute Engine and Amazon EC2. All tools and related services are maintained, and upon new developments, new modules will be added to the environment.

**Supplementary information:** Supplementary data are available at Bioinformatics online.

## 1 Introduction

Metabolomics has been increasingly used to discover new associations between metabolites and disease and investigate biochemical processes underlying disease activity. Liquid chromatography tandem mass spectrometry (LC-MS) affords accurate and comprehensive metabolite profiling measurements. High resolution, accurate mass-based methodologies are capable of measuring both metabolites of known identity, that are confirmed using authentic reference standards, and LC-MS peaks derived from yet to be identified metabolites that are tracked by measured retention time (RT) and mass to charge ratio (m/z). Resulting data sets are complex, containing thousands of LC-MS peak features measured across hundreds or thousands of samples. Before these datasets can be analyzed, it is often necessary to standardize the data to mitigate time-based or batch-based trends related to changes in instrument sensitivity that can occur over the course of data acquisition. Since a single metabolite can yield multiple ion adducts and ionized fragments in the mass spectrometer, redundant features must be identified and removed prior to analysis. Finally, it is necessary to acquire multiple batches of data over time for large studies of thousands of sample, but it is challenging to accurately "align" unknowns between batches due to the fact that there are many thousands of features and each display subtle differences in measured RT and m/z between batches. Our lab has created software tools to facilitate data processing and address these challenges. Here, we introduce *netome*, a computational suite that provides access to these data processing tools as well as software for interpreting data, such as testing their relationship with clinical data other profiling data such as microbial profiles. The reports include data and visualization for validation and interpretation.

## 2 netome framework

The *netome* framework is designed to provide tools and support for: 1) metabolite profiling, 2) statistical analysis and pattern discovery, and 3) computational platform and utilities. Prior to using *netome*, feature extraction tools such as Progenesis QI (Nonlinear Dynamics) or TraceFinder (Thermo Scientific) are used to batch-process raw LC-MS data files from individual samples into tabularized datasets consisting of feature information (such as measured masses, retention times, and metabolite names when known) and peak intensities (**Fig. 1a**). The *netome*’s metabolite profiling tools (**Fig. 1a)** use the tabularized raw data as input and generate processed data by applying a set of particular tasks including data scaling (*netomeScalar*), feature clustering (*neteomClust*, *mclust*), and multi-dataset alignment (*M2Aligner)* which combine profiles metabolites **Fig. 1b**. The output data table can be analyzed using statistical analysis and pattern discovery tools to identify significant metabolites and associations with other data types such as clinical phenotypes (*MaAsLin2,* tests metadata vs. data) and profiles such as microbiome profiles (*HAllA,* tests data vs. data, **Fig. 1c**). Computational platform and utilities provide an infrastructure for using and supporting these tools with extensive documentation and tutorials. The framework supports researchers to reproduce their analyses by providing a default setup for their runs and controlling of dependencies.

**Figure 1:**
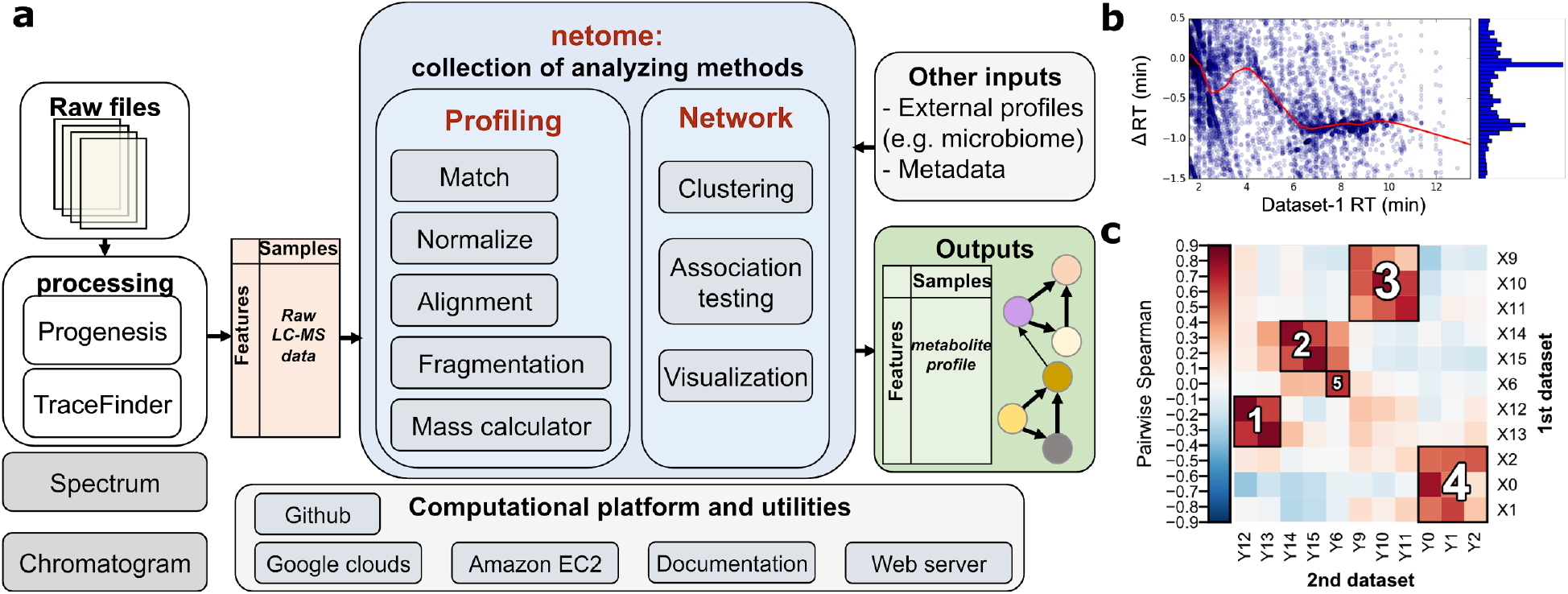
The netome framework: **a,** workflow overview. Tabularized, raw LC-MS data are inputs to *netome*. Profiling modules in *netome* produce a set of polished data and reports and network modules include statistical analysis and pattern discovery tools. **b,** metabolite profiling tools. *m2aligner* is a tool to combine two microbial profiles using an alignment technique. Here the plot shows the difference between metabolites retention time between two datasets. **c,** network tools. *HAllA*, an example of tools supported in *netome* for testing the potential association between metabolites (1st dataset) can be tested versus other biological profiles (2nd datasets) such as microbiome profiles.

## 3 netome package and installations

The *netome* package is an organizational structure and an interface to the tools available at https://broadinstitute.github.io/netome/. *netome café,* available at broad.io/netomecafe, as a part of the infrastructure of netome, is a web server to run tools online. The server has 1 Intel Haswell CPU, 3.75 GB RAM memory, and 30GB disk storage to support this service.

All the tools are supported by demonstration datasets and results with reports and visualization. The package can be scaled up to user desired machines on Google by using netome’s public Google Cloud image via the netome Google Cloud Bucket and Amazon EC2. We keep this image consistently update to reflect our new changes and added tools.

## 4 Conclusion

*netome* provides a framework to process and analyze metabolomics data and explore associations with other omics data and metadata. The computational package is available with extensive documentation and demonstrations.

## Acknowledgments

We appreciate the contribution of the metabolomics platform members and collaborators over many years to various netome’s modules. We also thank Broad’s COMM lab (Robert Majovski) for his feedback on the text. Amazon Web Services supported this work by an award to Ali Rahnavard for AWS Cloud Credits Research program.

## References

Clish, C. B. (2015). Metabolomics: an emerging but powerful tool for precision medicine. Molecular Case Studies, 1(1), a000588.

Rahnavard, G., et al. (2017). High-sensitivity pattern discovery in large multi’omic datasets. huttenhower.sph.harvard.edu/halla

Jang, C., Chen, L. and Rabinowitz, J.D., 2018. Metabolomics and Isotope Tracing. Cell, 173(4), pp.822-837.

Roberts, L.D., Souza, A.L., Gerszten, R.E. and Clish, C.B., 2012. Targeted metabolomics. Current protocols in molecular biology, 98(1), pp.30-2.

Vinayavekhin, N. and Saghatelian, A., 2010. Untargeted metabolomics. Current protocols in molecular biology, 90(1), pp.30-1.

Himel Mallick, et al. (2018). Multivariable Association in Population-scale Meta’omic Surveys. huttenhower.sph.harvard.edu/maaslin2

